# Variant U1 snRNAs contribute to cell cycle and differentiation control of human iPS cells

**DOI:** 10.1101/2025.09.14.676199

**Authors:** Yajie Zhu, Konstantinos Sofiadis, Milos Nikolic, Vasilisa Kalinkina, Athanasia Mizi, Lukas Cyganek, Carmelo Ferrai, Argyris Papantonis

## Abstract

Maintenance of stem cell identity as well as differentiation of stem cells into the various lineages requires intricate regulation of gene expression. Despite intensive research, our understanding of these regulatory networks remains incomplete. Here, we focus on the understudied paralogs of the U1 small nuclear RNA (snRNA) gene known as variant U1 (vU1) snRNAs. By generating isogenic knockout lines of human induced pluripotent stem cells for two different vU1s, we show that their loss profoundly changes both hiPSC gene expression and cell cycle profiles. These changes appear to manifest at the level of alternative splicing choices, including those at recursive splicing sites, and lead to the differential availability of stem cell regulators. Together, our results shed new light on the function of vU1 snRNAs and further our understanding of the programs controlling human pluripotency.

## INTRODUCTION

The maintenance of stem cell identity and the commitment of stem cells to a differentiation path both rely on the activation of specific gene regulatory networks^1^. Shifting from one to the other involves, amongst others, changes in cell cycle control, cell adhesion and morphology, metabolism, and protein availability^2–6^. Such changes can be achieved by the differential control of gene expression levels, as well as by changes to alternative splicing patterns. However, the molecular events and the key players in these changes are still not fully understood.

Transcription factors, epigenetic regulators and the ensuing control of transcription have been intensely studied in respect to stem cell maintenance and differentiation^7–9^, with alternative splicing and its regulation not receiving attention until much more recently^10,11^. Splicing choices are executed by ribonucleoprotein (RNP) complexes containing small nuclear RNAs (snRNAs) like U1, U2, U4-6 and their associated factors, the precise composition and dynamic functional states of which have been well described^12,13^. However, various animal genomes were found to carry additional copies of snRNA genes that were long considered pseudogenes^14–17^. Strikingly, most of these variant U-snRNA genes show highest expression during the earlier stages of development^18^, but their function remains enigmatic.

In humans, almost 1700 copies of U-snRNA gene variants have been identified^19^ (Vazquez-Arango & O’Reilly, 2018). They correspond to essentially all major (i.e., U1, U2, U4, U5, U6) and minor spliceosome snRNA-encoding genes (i.e., U11, U12, U4^ATAC^, U6^ATAC^) and although they have been regarded as pseudogenes, accumulating evidence argues in favor of their incorporation into active spliceosomal complexes and involvement in RNA processing^20–24^. For variant U1 genes, there exist >100 annotated copies on chr1^22.25^. When compared to the canonical U1 snRNA, various sequence differences, including base substitutions and deletions, arise, but only few overlap the Sm binding motif^19^. This can potentially alter their binding specificity along pre-mRNAs, whilst allowing formation of snRNP complexes with known spliceosome components^24,26^.

Human vU1 snRNAs were shown to be expressed highly in pluripotent or reprogrammed stem cells^19^, but their roles in respect to stem cell identity and functions remain unknown. To address this, we generated complete-knockout human induced pluripotent stem cell (hiPSC) lines for two different vU1 genes and used genomics and functional assays to investigate their effects on hiPSC homeostasis. This way, we were able to show that their loss-of-function produces pronounced gene expression and cell cycle changes that can be linked to perturbed alternative splicing patterns genome-wide.

## RESULTS

### vU1 snRNA ablation affects hiPSC cell cycle progression

Human vU1 snRNA copies on chr1 (**Fig 1a**) are expressed to different extents in different cell types and developmental stages. We decided to focus on two such variant genes, vU1.3 and vU1.8, for two reasons. First, because they are markedly more expressed in embryonic tissues and organs, and overall up to ∼20% the levels of canonical U1 snRNA (**Fig S1a**). The expression of the two variant U1s correlated highly with one another in the tissues queried, but not with the expression of canonical U1 (**Fig S1b**), which might hint to non-overlapping functions of the resulting snRNPs. Second, because of their deviation in sequence relative to the canonical U1. vU1.3 carries a C-to-T mutation at its 5’ splice site recognition domain, which could allow it to recognize AT instead of canonical GT splice donor sites (**Fig S1c**). vU1.8 carries an almost intact 5’ splice site recognition domain and a small deletion at its U1-A protein binding site^22^. Both these vU1 genes also carry multiple other base substitutions, including some in their U1-70K binding domain (**Fig S1c**), and overall deviate enough so that specific gRNAs could be designed for their genetic ablation (see **Table S1**).

**Fig 1.**
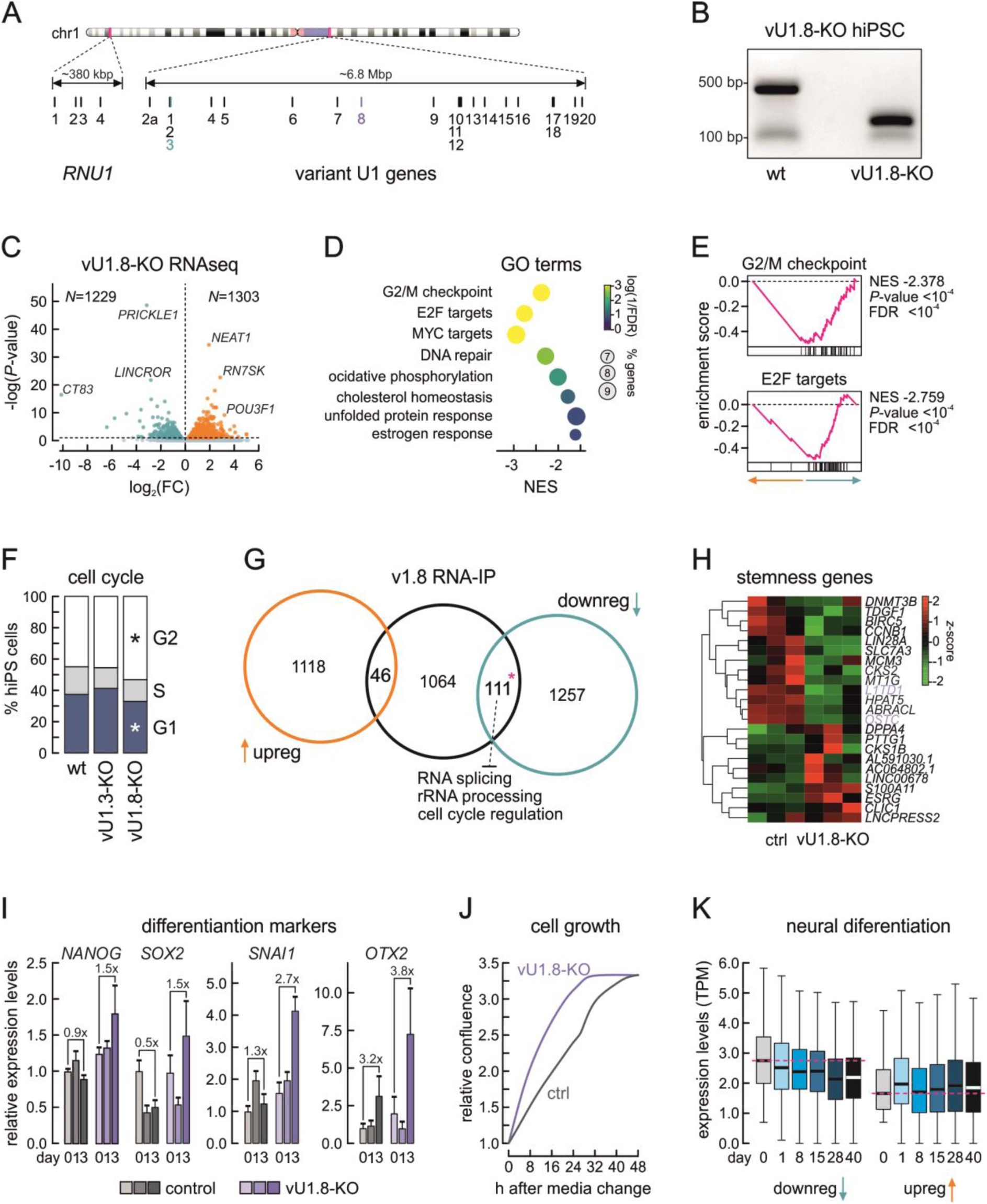
The vU1.8 knockout affects gene expression and the cell cycle in hiPSCs. **a**, Ideogram of human chr1 (top) with the regions harboring canonical (bottom left) and variant U1 snRNA genes detailed (bottom right). The *RNU1*-3 (green) and *RNU1*-8 loci (purple) are indicated. **b**, Electrophoretic profiles of PCR products corresponding to the wild-type (wt) and knocked-out RNU1-8 locus (vU1.8-KO). Molecular weight marker positions are indicated. **c**, Volcano plot showing differentially up-(orange) and downregulated genes (green) in vU1.8-KO cells given a *P*_adj_ cutoff of <0.05. **d**, Plot showing the top gene ontology (GO) terms associated with vU1.8-KO downregulated genes from panel c. **e**, Plots showing GSEA results for with vU1.8-KO downregulated genes from panel c. **f**, Bar plots showing the percentage of cells in each cell cycle phase. \**P*<0.05, Fisher’s exact test. **g**, Venn diagram showing the overlap of differentially expressed genes from panel c with transcripts co-immunoprecipitating with vU1.8. \**P*<0.05, hypergeometric test. **h**, Heat map showing expression of known stemness-related genes in control (ctrl) and knockout hiPSCs (vU1.8-KO). Genes directly bound by vU1.8 are highlighted (purple). **i**, Bar plots showing changes (relative to day 0 ±S.D.) in expression levels of the indicated genes for control (ctrl) and knockout hiPSCs (vU1.8-KO) at 1 or 2 days of differentiation. **j**, Line plots showing changes in relative control (ctrl) and knockout hiPSCs (vU1.8-KO) confluence in the 48 h after initiating differentiation. **k**, Box plots showing the expected changes in expression levels for the 1229 down- (left) and the 1303 upregulated genes during the differentiation of hiPSCs into neuronal organoids (from ref. ^29^).

Using CRISPR/Cas9-mediated knockouts, we managed to delete all vU1.3 or vU1.8 copies in hiPSCs and generate multiple single cell-derived vU1.3-KO and vU1.8-KO clones (**Figs 1b** and **S2a**). Knockout lines did not exhibit apparent phenotypic differences to wild-type hiPSCs or to one another (**Fig S2b**). However, each vU1 knockout had pronounced and non-overlapping effects to hiPSC gene expression patterns, examined by RNA-seq on at least two independent knockout clones for each genotype. Ablation of vU1.8 produced the strongest effects, with ∼1300 up- and >1200 down-regulated genes (given an adjusted *P*-value cutoff of 0.05; **Fig 1c**). Among the genes transcriptionally suppressed were those encoding the key “reprogramming” transcription factor genes *MYC* and *SOX2* ^27^, indicating perturbed stem cell identity maintenance. More broadly, downregulated genes were primarily associated with gene ontology (GO) terms related to cell cycle regulation (i.e. “G2/M checkpoint”, “E2F targets”, “MYC targets”) in gene set enrichment analyses (**Fig 1d,e**). Accordingly, cell cycle profiling of PI-stained control (maternal) and vU1.8-KO hiPSCs revealed an accumulation of cells in the G2 phase of the cell cycle at the expense of G1/S-phase cells in knockout clones; this profile differs significantly from that of wild-type cells (**Fig 1f**).

However, this was not the case for vU1.3-KO cells. Here, we found an order of magnitude fewer differentially expressed genes in knockout cells, with 122 up- and 244 down-regulated (given the same cutoff as above; **Fig S2c**). GO term analysis showed highest enrichment for terms like “DNA conformation change” and “RNA metabolism” in relation to upregulated genes, whereas downregulated ones were associated with “wound response”, “covalent chromatin modification”, “chromatin remodeling”, “growth stimulus response” (**Fig S2d**). Despite the apparent differences in the magnitude and type of gene expression changes triggered, the two vU1 knockouts converge in that they both deregulate genes involved in nucleosome assembly, RNA metabolism and splicing, as well as in Wnt and Notch signaling (see **Table S2**) that are important for hiPSC differentiation.

As is usually the case, generating knockout lines comes with both direct and indirect (or compensatory) effects to gene expression. To discriminate between them, we immuno- precipitated RNAs interacting with either the vU1.8 or the vU1.3 snRNA using the knockout lines as baseline controls. These IPs retrieved >1400 RNAs interacting with vU1.3 and >1200 interacting with vU1.8, of which 203 were shared (**Fig S2e**). Interestingly, the RNAs in both IPs were on average significantly longer than those in the control IP (**Fig S2f**) and contained many genes encoding cell cycle regulators, chromatin assembly and remodeling factors (e.g. *CCNB1*, *CCNA2*, *SMC3*, *SMARCA1/-C1/-D1* or *CHAF1A*; **Table S3**).

We crossed the lists of differentially regulated genes (DEGs) from each knockout with these RNA-IP catalogues to find that only ∼10% of vU1.3-KO DEGs were direct targets of the vU1.3 snRNAs (**Fig S2g**). For vU1.8, this percentage further drops to ∼4% for the up- and to 8.5% for the down-regulated genes in vU1.8-KO hiPSCs (**Fig 1g**). Nevertheless, the 25 down- regulated vU1.3 targets and the 111 of vU1.8 are more than expected by chance, and they are enriched for genes associated with histone modifications, lipid transport, and FGF signaling for vU1.3 (**Fig S2G**) and with RNA splicing, rRNA processing and cell cycle control (**Fig 1g**). For vU1.3, we also managed to retrieve a protein interactome using the same IP approach. Here, we catalogued 113 robust protein-vU1.3 interactions that were associated with GO terms like “RNA splicing”, “mRNA processing”, “RNA localization/stabilization” and “RNP assembly”, as well as “DNA conformation” and “rRNA maturation” (**Fig S2h**). Notably, this interactome showed minimal overlap with the known protein interactomes of the canonical U1 and U11 snRNAs (**Fig S2i**), although vU1s have been shown to incorporate into functional RNPs^24^ and that the term “minor spliceosome” is significantly enriched for vU1.3 interactors (**Fig S2h**).

### vU1.8, but not vU1.3 ablation perturbs hiPSC transcriptional homeostasis

The pluripotent nature of hiPSCs allows them to give rise to all cell types in the three major developmental lineages. We therefore wanted to ask whether either vU1 knockout affects genes linked to stem cell identity maintenance or to the ability of hiPSCs to differentiate. We based this question on our observation of higher vU1 snRNA expression in embryonic tissues (**Fig S1a**), and on data showing vU1 upregulation during cell reprogramming into iPSCs^19^.

To this end, we queried the expression of 23 genes considered as “stemness” markers in hiPSCs^28^. 13 of these were consistently downregulated in vU1.8-KO cells, while 4 consistently upregulated (**Fig 1h**). Of the former 13 genes, *L1TD1* and *OSTC* were also directly bound by vU1.8 in RNA-IP experiments (**Table S3**). In vU1.3-KO cells, such changes were only detected for two genes, *DNMT3B* and *SLC7A3* (not shown). Next, we tested the response of knockout hiPSCs to non-directed differentiation (via media replacement; see **Methods**). We measured changes in expression at 1 and 2 days after differentiation was induced to find that key stem cell markers like *NANOG* and *SOX2* were inducedby ∼1.5-fold at day 2 in the absence of vU1.8, along differentiation markers like *OTX2* and *SNAI1* that were also over-induced compared to control cells (**Fig 1i**). Again, such effects could not be observed in vU1.3-KO cells (**Fig S2j**). Notably, these gene expression changes in vU1.8-KO cells were accompanied by higher growth rates compared to control hiPSCs (**Fig 1j**), which is in line with our RNA-seq data analyses (**Figure 1c-f**).

Last, since the knockout of vU1.8 led to the deregulation of stemness genes (**Fig 1h**) and of TFs involved in neural development (e.g., OTX2; **Fig 1i**), we examined how the levels of vU1.8-KO DEGs compare against transcriptional changes occurring during the generation of neuronal organoid generation from hiPSCs^29^. Looking at RNA-seq data from 0 to 40 days post-differentiation, we found that genes suppressed in the absence of vU1.8 would normally be most downregulated in later neurogenesis stages (days 28-40; **Fig 1k**). On the other hand, gene already induced in vU1.8-KO cells would normally be upregulated within 1 day of differentiation and then again in much later stages (day 28; **Fig 1k**). These analyses suggest that vU1.8-KO hiPSCs exhibit perturbed transcriptional profiles that, in part, liken those invoked in later development.

### vU1 ablation leads to widespread alternative splicing changes in hiPSCs

As U1 snRNAs are essential spliceosome components involved in 5’ splice site recognition, we sought to assess the contribution of vU1.3 and vU1.8 to alternative splicing (AS) choices. AS in stem cells can lead to inclusion or removal of protein domains encoded by mRNAs, or affect their subcellular localization, coding potential and stability^30^. Thus, we first used IsoformSwitchAnalyzeR^31^ in order to annotate and quantify the most pronounced changes in mRNA isoforms in each vU1-KO hiPSC line. Of the hundreds of isoform switch events detected upon knockout of each vU1 snRNA, only 6 are shared between the two KOs – but show enrichment for genes in important pathways like embryonic development, tube morphogenesis or p53 signaling. vU1.3-KO hiPSCs show a significant number of isoform changes leading to a loss of encoded functional domains (*N*=92) and to shortening of open reading frames (ORFs, *N*=59; **Fig 2a,b**). On the other hand, vU1.8-KO cells only show a significant number of changes (*N*=130) when it comes to loss of ORFs from mRNAs (**Fig 2a,b**). For example, *ZNF774*, a gene suppressor of Notch signaling^32^, and *SHC4*, a signaling modulator in stem cells^33^, both show an upregulation of non-coding isoforms at the expense of mRNAs with intact ORFs upon vU1.3- and vU1.8-KO, respectively (**Fig S3a**).

**Fig 2.**
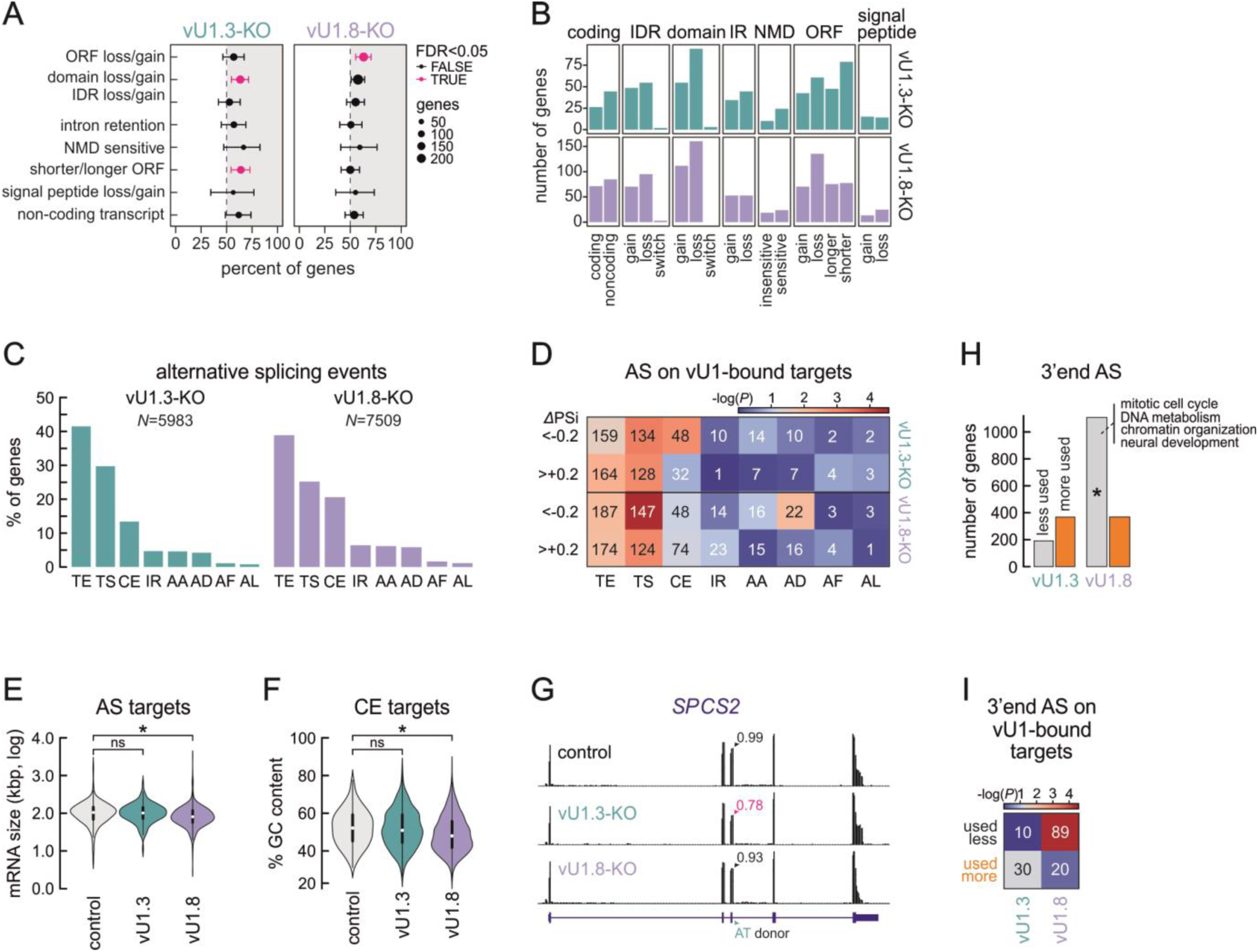
Effects on alternative splicing choices in vU1-KO hiPSCs. **a**, Plot showing the percent of all active genes that show any of the eight mRNA isoform consequences indicated due to alternative splicing (AS) in each vU1-KO hiPSC line. Isoform changes that occur significantly more (FDR<0.05) are indicated (magenta). **b**, Bar plots showing the number of genes that show any of the indicated mRNA isoform changes in vU1.3- (top) or vU1.8-KO hiPSCs (bottom). **c**, Bar plots showing the percentage of genes exhibiting at least one of the indicated AS changes in vU1.3- (top) or vU1.8-KO hiPSCs (bottom). The total number of genes (*N*) with AS changes is indicated. TE, alternative transcription end site; TS, alternative transcription start site; CE, alternative core exon usage; IR, intron retention; AA, alternative acceptor site; AD, alternative donor site; AF, alternative first exon; AL, alternative last exon. **d**, Heat map showing the number of events per AS type for vU1-associated transcripts in vU1.3- (top) or vU1.8-KO hiPSCs (bottom). The color code represents statistical overrepresentation (-logP) in each case. **e**, Violin plots showing the distribution of lengths for all alternatively spliced mRNAs (grey) or for vU1.3- (green) and vU1.8-bound transcripts (purple). *: *P*<0.01, Wilcoxon-Mann-Whitney test. **f**, As in panel e, but for the mean GC content of these mRNAs. **g**, Exemplary genome browser views of RNA-seq coverage along the *SPCS2* locus from control (top), vU1.3-KO (middle), and vU-1.8-KO hiPSCs (bottom). The percent usage of the indicated AT donor site is shown. **h**, Bar plots showing the number of genes with reduced (grey) or increased alternative 3’ end usage (orange) in vU1.3-KO (left), and vU-1.8-KO hiPSCs (right). *: *P*<0.01, Fisher’s exact test. The four most enriched GO terms for the mRNAs showing reduced 3’ end AS are shown. **i**, As in panel d, but for alternative 3’ end usage in each knockout line.

To obtain a more precise understanding of the AS changes induced by each vU1-KO, we used Whippet^34^ to map and quantify individual AS events from RNA-seq data. This way, we mapped ∼6000 and >7500 AS events in vU1.3- and vU1.8-KO hiPSCs, respectively. Of these, ∼40% concerned alternative 3’ end (TE) and 25-30% alternative transcription start (TS) site usage in both vU1 knockouts (**Fig 2c**). In vU1.8-KO hiPSCs, there were also >20% of events involving the alternative inclusion/skipping of core exons (CE; **Fig 2c**). Still, it was not clear how many of these events were a direct consequence of vU1 loss, since each knockout also induced or suppressed the expression of various splicing factors (**Table S2**). We therefore crossed the list of AS events (**Table S4**) with the mRNA targets of each vU1 snRNA (**Table S3**). This narrowed down the list of AS mRNAs to ∼700 for vU1.3- and to >860 for vU1.8-KO hiPSCs, but did not change the fact that TS and TE events were significantly overrepresented in both cases (**Fig 2d**). Curiously, and despite the fact that the vU1 mRNA targets were longer than average (**Fig S2f**), direct vU1.8 AS-targets were shorter and of lower GC content compared to control mRNAs (**Fig 2e,f**).

As alternative TE events were the most prevalent in either vU1-KO line, and the canonical U1 snRNP has been shown to suppress premature 3’-end cleavage and polyadenylation via “telescripting”^35^, we analyzed 3’ end AS events further by quantifying 3’ UTR usage in respect to the levels of the last exon in each mRNA^36^. This analysis showed that the knockout of vU1.8 leads to significantly more 3’ UTR shortening and down-regulation compared to both control and vU1.3-KO hiPSCs (**Fig 3h**). GO term analysis also showed that genes with shortened 3’ UTRs and/or less usage in vU1.8-KO cells were associated to cell cycle, DNA metabolism and chromatin organization control, as well as to neuronal differentiation (**Fig 3h**). One such example would be *TIMP2*, encoding a factor involved in ECM degradation that was shown to promote hPSC renewal^37^. Next, we looked for direct targets by considering the RNA-IP catalogues from each vU1 (**Figs 1g** and **S2g**). This confirmed the strong enrichment for shorter and less used 3’ UTRs in vU1.8-KO hiPSCs, with 89 mRNAs being regulated this way (**Fig 3i**).

**Fig 3.**
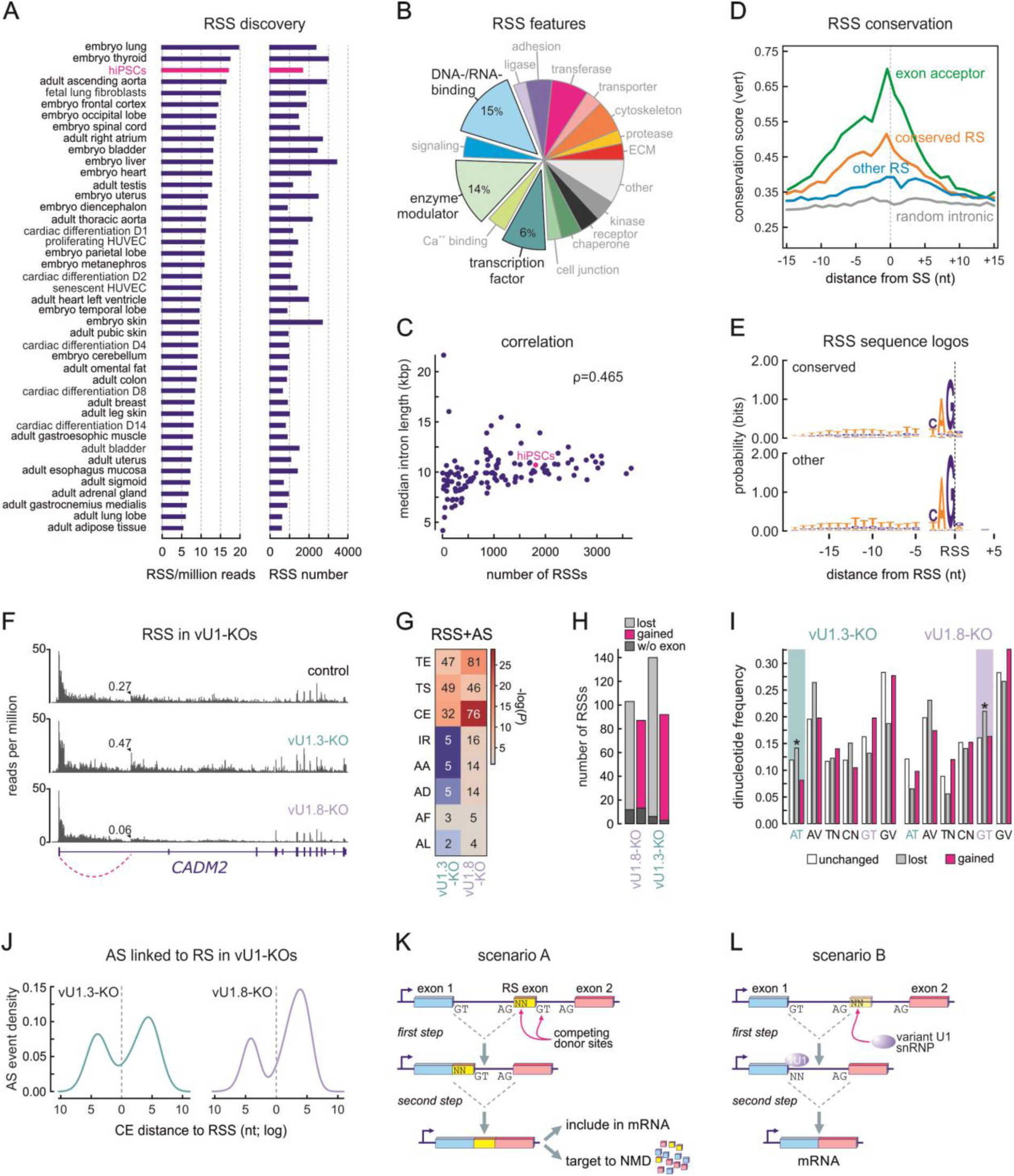
vU1 contribution to recursive splicing in human cells. **a**, Bar plot showing the number of recursive splicing sites (RSSs) per million RNA-seq (left) and their total number per each publicly available dataset. **b**, Pie chart showing the percent of genes carrying RSSs from panel a that fall into different biological function GO terms. The three top categories are highlighted. **c**, Plot showing the correlation between total RSS numbers and median intron length per each cell type from panel a. The overall Pearson’s correlation coefficient is indicated. **d**, Line plot showing the conservation score (from all vertebrates in the UCSC browser) in the 30 nt around canonical 3’ acceptor sites (green), RSSs with conservation score >0.45 (orange) or <0.45 (blue), and randomly selected sites from the same introns (grey). **e**, Logos showing the sequence probability at each position around conserved (score >0.45) and all other RSSs (score <0.45) from the data in panel a. **f**, Genome browser views of total RNA-seq from the CADM2 locus showing alternative usage of an exemplary RSS (arrowheads) in control (top), vU1.3-KO (middle), and vU1.8-KO hiPSCs (bottom). **g**, Heat map showing the number of RS Ss that are also linked to AS events in vU1.3- (left) or vU1.8- KO hiPSCs (right). The color code represents statistical overrepresentation (-log*P*) in each case. **h**, Bar plots showing the number of RSSs lost (grey) or gained (magenta) in vU1.3- (left) and vU1.8-KO cells (right) compared to control hiPSCs. The number of RSSs not associated with an immediate downstream exon are indicated for each case (dark grey). **i**, As in panel h, but for specific RS dinucleotides (N: A, T, C, or G; V: A, C, or G). The RS donor dinucleotides matching vU1.3 (left) and vU1.8 sequences (right) are highlighted. \**P*<0.05, Fischer’s exact test. **j**, Plots showing the density of exon ends up- and downstream of RSSs in vU1.3- (left) and vU1.8-KO hiPSCs (right). **k**, A model for how RSS-exon inclusion in mRNA via competition between RS and canonical donor sites may lead to the production of translatable or NMD-prone transcripts. **l**, As in panel k, but for RSSs not followed by a cryptic exon.

Another notable finding was that, in these vU1.3 AS targets, alternative CE usage was significantly overrepresented (**Fig 2d**) in contrast to what we observed in the whole AS dataset (**Fig 2c**). One such example is exon 3 of *SPCS2*, a U12-type cassette exonthat contains an AT 5’ splice site. This exon is included >20% less in mRNA in vU1.3-, but not in vU1.3-KO hiPSCs (**Fig 2g**). This matches the fact that vU1.3 has a C-to-T mutation in its 5’SS recognition site, theoretically enabling it to recognize AT donor dinucleotides, while vU1.8 does not (**Fig S1c**). At the same time, vU1.8 AS targets showed significant enrichment for the skipping of alternative donor sites (AD; **Fig 2d**). Among the 22 AD vU.18 targets are *CPSF7* — a key player in pre-mRNA cleavage and polyadenylation^38^, *ARPP19* and *CCDC57* — both regulators of the G2/M cell cycle transition^39,40^, and *BAZ2A* and *SMARCA2* — components of chromatin remodeling complexes linked to stem cell maintenance and proliferation^41–43^.

Last, we examined the extent to which AS changes were coupled to changes in expression levels. Intersecting vU1-KO DEGs (**Figs 1c** and **S2c**) with genes undergoing AS, we found significant overrepresentation only in vU1.8-KO cells and only for CE inclusion and intron retention (**Fig S3b**). While both increased and decreased intron retention correlated with upregulated gene expression, it was CE inclusion (but not skipping) that predominantly correlated with higher expression levels (**Fig S3c**). This latter effect could be explained by the mechanism of “exon-mediated activation of transcription starts”^44^, whereby exon inclusion in mRNAs often activated transcription from exon-proximal weak upstream promoters. In fact, when comparing the distribution of CEs related to annotated gene transcription start sites (TSSs), we found that those included more in the transcripts of vU1.8-KO upregulated genes were located significantly closer to their respective TSS than all other CEs in DEGs (**Fig S3d**). Similarly, alternative 3’ UTR usage in vU1.8-, but not in vU1.3- KO hiPSCs was found linked to differential gene expression. 134 mRNAs showing decreased usage of an alternate 3’ UTR were significantly downregulated (e.g., *TOP2A* that encodes a key topoisomerase for cell division^45^), and the converse applied to 46 mRNAs with decreased usage of an alternate 3’UTR (e.g., *CTNNB1* that encodes for β-catenin^46^). Notably, the latter were enriched for genes under the control of transcription factors known to regulate cell cycle progression, like p53, MYC and E2F1 (**Fig S3e**). This aligns well with the cell cycle effects observed for vU1.8-KO cells.

### Recursive splicing patterns are altered in the absence of vU1 snRNPs

Studies in various organisms, including in fruit flies^47,48^ and man^49,50^, have described an additional layer in splicing, whereby (mostly longer) introns may be removed in a stepwise manner rather than via a single reaction. This is accordingly called “recursive splicing” (RS) and involves the splicing of 5’ donor dinucleotides into cryptic sites located deep inside of introns (reviewed in refs^51,52^). In flies, this occurs at YAG|GU “zero-length” exons^47^. In man, however, we previously documented intronic sites producing RS intermediates that did not always carry the conventional GU sequence. Here, the resulting RS donor was not a GN dinucleotide in >55% of cases^49^. Given that vU1 snRNAs have 5’ splice site binding sites that deviate from the canonical one, we asked whether they might also be involved in the recognition of specific RS site subsets in hiPSCs.

To address this question, we first mapped hybrid reads representing RS events (i.e., 5’ splice donors at the end of exons spliced into YAG|NN intronic sequences^49^) across a collection of ENCODE and Epigenome Roadmap total RNA-seq data. We again found that RS is on average more prevalent in cell types of embryonic origin (**Fig 3a**). As many of these might be non-productive splicing events (discussed in ref.^52^), we sought to obtain some validation data. First, we exploited ENCODE eCLIP data^53^ to assess the enrichment of RNA-binding proteins (RBPs) at RS sites. Indeed, we discovered that a subset of RBPs, e.g. LARP4, the helicases DDX5/-6 and IGF2BP1/-3, all bind around RSS withs characteristic double-peak profiles (**Fig S4a**). Second, we confirmed the existence of 8 of these RS intermediates that did not involve a canonical GT in their RS sequence in HEK293 cells (**Fig S4b**), and then used CRISPR/Cas9 genome editing to specifically microdelete two different ones and assess their contribution to mRNA production from their host gene. Here, microdeletion of the RS site in the first intron of the *UBE2E2* gene led to a >50% reduction in mRNA levels, whereas that in the first intron of *UQCC1* had no effect (**Fig S4b**). This is in line with our previous testing of different RS sites within the first long intron of *SAMD4A* in human endothelial cells^49^. Next, we looked into the attributes of RS events across cell types. Collectively, there appears to be a bias for RS occurring in genes encoding DNA- and RNA-binding proteins, modulators of enzymes or transcription factors (**Fig 4b**). RS occurrence was not exclusive to long introns, but their numbers did scale with the median length of the intron in which they occur (**Fig 4c**). We also stratified RS sites into conserved (*phastcons* score >0.45 across vertebrates) and non-conserved ones (score <0.45; **Fig 4d**), but found essentially no bias in sequence composition between the two (**Fig 4e**), which argues against sites with canonical RS donors being more conserved (and presumably functionally relevant).

Finally, we hypothesized that if RS sites are indeed functionally relevant, they may contain single nucleotide polymorphisms (SNPs) linked to diseases or traits in a cell type-specific manner. Based on the idea that 70-90% of SNPs associated with common complex diseases and traits occur in non-coding parts of the genome, including introns, we had previously developed GARLIC, a software for identifying potentially disease-causative genetic variants that overlap regulatory sequences of interest^54^.

In brief, GARLIC is linked to a database of SNPs drawing from genome-wide association studies and we used it to assess any statistical enrichment of genetic variants in the 25 nt around RS sites (i.e., 20 nt up- and 5 nt downstream of the RSS to also cover its associated T-tract). Initially, we used a catalogue of RSSs from a time course of hiPSC differentiation into cardiomyocytes^55^ along which we saw a gradual decrease of RS events (**Fig S4c**). However, SNPs associated with cardiomyopathy and echocardiogram traits were found specifically enriched in cardiomyocyte RS sites (and not in earlier progenitors; **Fig S4c**). Thus, we next expanded our tests to the whole GARLIC database using RS catalogues from all available cell types (see **Fig 3a**). This revealed various clusters of relevant cell types and traits, like RS sites from neuronal cell types being enriched for SNPs associated with migraines, bipolar and neurodegenerative disorders, psychosis, and depression (**Fig S4d**). Our analysis therefore provides additional indications that RS is likely consequential for gene regulation.

Given the pervasive presence of non-GT donors at RS sites, we asked whether vU1 snRNAs can be implicated in RS implementation. hiPSCs, in which vU1s are most expressed (**Fig S1a**), show one of the highest RS occurrences (**Fig 3a**). Thus, we applied additional filtering criteria for RS events (adapted from ref.^56^) and quantified RS usage by normalizing junction reads including an RS site to all junction reads that include their upstream exon. This way, we can statistically determine the over- or underrepresentation of a given RS event in vU1-KO cells compared to control hiPSCs (see example in **Fig 3f**). In total, we identified ∼200 and >230 differentially used RS sites in vU1.3- and vU1.8-KO cells, respectively. In both knockout lines, more RSSs were lost than gained (**Fig 3g**), and the genes carrying gained or lost RSSs in vU1.8-KO cells were enriched for pathways linked to early development and cell differentiation.

To test whether differentially used RSSs correlate with alternative splicing, we crossed the two datasets for each knockout hiPSC line. This showed that vU1.3 ablation may be equally correlated with alternative usage of transcription start and end sites, as well as with core exon usage, whereas vU1.8 loss produces a similar profile but with a strong enrichment for differential CE usage (**Fig 3h**). In fact, when it comes to the location of these alternatively included/skipped CEs, we found that the gain or loss of an RS event predominantly affect exons located downstream of the RS site, directly implicating recursive splicing in mRNA composition outcomes (**Fig 3i**).

We next tested whether hiPSC RS sites were “zero-length” exons (as in Drosophila^48^) or involved an RS-exon (as proposed for human neuronal cells^50^). RS-exons are thought to provide “exon definition” to RS sites, but not be included in mature transcripts^50^. In our hiPSC lines, the vast majority of RS sites (89% or more) were followed by an RS-exon (**Fig 3g**). However, a non-negligible number of these RS-exons display signal enrichment in poly(A)-selected RNA-seq data from hiPSCs, which means that they are likely cryptic alternative exons, the usage of which is under the control of vU1.3 or vU1.8 (see **Fig S3b**).

Finally, we asked to which extent the loss of vU1.3 or vU1.8 would lead to selective loss or gain of RS usage based on the RS-donor motif that the variant snRNAs might recognise. In the case of the vU1.3-KO, we recorded a significant decrease in the usage of RSSs with an AT donor with a concomitant increase in GN usage (**Fig 3j**). Conversely, in vU1.8-KO cells, we recorded a significant drop in GT RS-donor usage, but with a concomitant increase in GV ones (V=A/C/G; **Fig 3f,j**). These effects are consistent with the theoretical ability of vU1.3 to recognise AT dinucleotides (due to a C-to-T mutation in its 5’ ss recognition site), which is not the case for vU1.8 that carries a canonical sequence (**Fig S1c**). Notably, we saw no bias in the differential loss or gain of conserved (phastcons score >0.45 across vertebrates) over non-conserved RS sites (score <0.45) in either vU1 knockout. Taken together, our analyses suggest that vU1.3 and vU1.8 do affect RS patterns genome-wide in specific gene subsets.

## Discussion

Even though variant U1 snRNAs are most expressed in human stem cells, their expression quickly declining upon differentiation^18^, their functional roles in gene expression control remained understudied. This was due to both the predominance of canonical U1 snRNA expression (vU1.3 and vU1.8 express at <1/3 the U1 levels in the cell types we looked at; **Fig S1a,b**) and the presumed redundancy among the many vU1 gene copies (see **Fig 1a**). We therefore selected two such genes, *RNU1-3* and *RNU1-8*, which could be specifically deleted from chr1 of hiPSCs in order to study the consequences of their loss on stem cell gene regulation and identity maintenance.

Using these hiPSC knockout lines, we found that the ablation of either vU1 snRNA leads to profound gene expression changes, albeit to different extents (vU1.8-KO have one order of magnitude more DEGs compared to vU1.3-KO; **Figs 1c** and **S2c**). Most of these effects on gene expression appear to be indirect, as they only concern a subset of vU1 target RNAs (<13% and <3% for vU1.8- and vU1.3-KO targets, respectively, as determined using RNA-IPs; **Figs 1g** and **S2g**). At the same time though, and as might be expected for snRNAs that can assemble into apparently functional snRNPs^24^, both knockouts produce strong changes in hiPSC splicing patterns, with hundreds of their targets exhibiting alternative usage of transcription start and end sites, and of core exons (**Fig 2d**). This finding corrects previous postulations (based on mini-gene reporters) of limited vU1 contribution to splicing^57^. We then asked what the implications of these changes to hiPSC homeostasis are. These seem to primarily relate to the vU1.8 rather than the vU1.3-KO, likely reflecting their difference in effect magnitude. First, and despite the fact that no discernible phenotypic changes were seen, the loss of vU1.8 results in cell cycle effects, with KO-hiPSCs exhibiting a shorter G1 phase and faster growth (**Fig 2f,j**). This can, for example, be in part explained by the *MYC* transcript being a direct vU1.8 target that is both downregulated (log_2_FC=-0.7) and alternatively spliced at its two putative TSSs upon knockout. Second, in the absence of vU1.8, key transcription factor genes like *SNAI1* and *OTX2* showed aberrant expression patterns upon non-directed differentiation (**Fig 2i**). This agrees with the broad and early changes in the expression of many genes known to be differentially regulated during a neuronal differentiation time course (**Fig 2k**), including genes in the critical Wnt and Notch pathways. Interestingly, there exist reports linking the deletion of the chromosome region (1q12-21) that harbors the vU1 snRNA gene cluster to severe pediatric neurological dysfunctions^58^, while motor neuron cultures derived from SMA patients sustained abnormally high vU1 expression^19^. Both these examples further link vU1 regulation to proper developmental progression.

Finally, we entertained the hypothesis that vU1 snRNAs, due to their potential for pairing to variable 5’ splice site dinucleotides, might be also involved in the regulation of recursive splicing events^52^. Here, we did record a bias in the usage of AT or of GT RSS 5’ splice site donors in vU1.3- and vU1.8-KO cells, respectively (**Fig 3i**), despite the fact that we expect high redundancy in RSS site recognition and usage^49^. Interestingly, the absence of either vU1 studied here led to changes in RSS patterns that could be linked to the alternative splicing of the relevant transcripts (**Fig 3g**). This might be in turn connected to the fact that the majority of RS sites are followed by small cryptic RS-exons. These exons are likely competing with for inclusion into mRNA with their flanking exons, but many include sequences that target them for degradation (as has been proposed for neuronal cells^50^). Our data suggest that vU1 binding to these RS sites can signal the skipping of such RS-exons, providing another layer of RNA quality control (**Fig 3k,l**). Still, a non-negligible number of RS events likely contributes to the stepwise removal of introns and, thus, to the production of mature coding transcripts. This, alongside the various traits of RS-containing transcripts (**Figs 3b** and **S4a-d**), supports a functional role for this regulatory layer of splicing in human biology and disease.

Taken together, this work assigns vU1 snRNAs with a far more decisive role in human stem cell function and maintenance that previously appreciated. Nevertheless, this now begs a number of new questions. For example, long-read sequencing can be used to investigate the coincidence or not of many of these vU1-dependent alternative splicing effects on single transcripts. Similarly, despite having catalogues of vU1.3 and vU1.8 target transcripts, we are still lacking high-resolution information of their exact positions of interaction on RNA. Last, more functional testing of the contribution of RS in mRNA production is needed, as is a broader screen of the precise contribution of other vU1s to the RS phenomenon.

## Methods

### Cell culture

Human induced pluripotent stem cells (hiPSCs; GM24581*B from Coriell) were derived from the preprogramming of human GM02036 fibroblasts by overexpressing the four Yamanaka factors (i.e., hOCT3/4, hSOX2, hKLF4 and hL-MYC) via episomal vectors, and validated for genomic integrity and their pluripotent character. hiPSCs were grown in StemFlex media and dissociated using accutase (Sigma-Aldrich) at 37°C for 10 min when confluent. For non-directed differentiation, hiPSCs were seeded on Matrigel-coated plates in StemFlex Medium with 2 μM Thiazovivin at a density of 4.7×10^4^ cells/cm^2^ for confluence analysis or 2.1×10^4^ cells/cm^2^ for RNA extraction. The following day, the medium was replaced by DMEM (Gibco) supplemented with 10% FBS (Capricorn Scientific,) and 2 mM Glutamine (Thermo Fisher) and cultured for 1 to 3 days, with daily medium changes. For confluence analysis, plates were imaged every 30 min for 48 hours following differentiation induction in an IncuCyte platform (Sartorius), and confluence was quantified using the Incucyte AI Confluence Analysis™ workflow.

### CRISPR/Cas9 gene knockout

For generating vU1-knockout iPSCs, WT cells were genome-edited using ribonucleoprotein-based CRISPR/Cas9 with crRNA/tracrRNA and Hifi SpCas9, targeting upstream and downstream of the locus via tailored gRNAs (see **Table S1**). 300 pmol of Alt-R CRISPR-Cas9 crRNA and 300 pmol of Alt-R CRISPR-Cas9 tracrRNA were pre-assembled with 122 pmol Alt-R Hifi SpCas9 Nuclease 3NLS (all IDT DNA Technologies) to form ribonucleoprotein complexes, and were nucleofected in to 2×10^6^ early-passage (p<20) iPSCs using the 4D Amaxa Nucleofector system (Lonza; program CA-137) and the P3 Primary Cell 4D-Nucleofector X Kit (Lonza) according to the manufacturer’s instructions. Following nucleofection, iPSCs were replated into a Matrigel-coated (growth factor-reduced, BD Biosciences) 6-well plate containing StemFlex medium supplemented with 2 µM thiazovivin (Merck) and 100 U/ml penicillin and 100 µg/ml streptomycin (Thermo Fisher). After 3 days, transfected iPSCs were singularized using the CellenOne dispenser (Cellenion/Scienion) in StemFlex medium into Matrigel-coated 96-well plates. Successful genome editing was identified by Sanger sequencing and several homozygous knockout iPSC lines were established. Generated iPSC lines were maintained on Matrigel-coated plates, passaged every 4-6 days with Versene solution (Thermo Fisher) and cultured in StemMACS iPS-Brew XF medium (Miltenyi) supplemented with 2 µM thiazovivin on the first day after passaging with daily medium changes.

### Reverse-transcription quantitative PCR analysis

Total RNA from control and differentiated hiPSCs was treated with TURBO DNase (Invitrogen) to remove genomic DNA contaminants, and reversed-transcribed using the SuperScript™ IV Reverse Transcriptase (Invitrogen) protocol and random hexamers in the presence of RiboLock RNase Inhibitor (Thermo) in each reaction. The resulting cDNA was used in real-time quantitative PCR (qPCR) reactions using the qPCRBIO SyGreen® Mix Separate-ROX system (PCR Biosystems) as per the manufacturer’s instructions in 384-well PCR plates and a final reaction volume of 10 μL. A qTower 384G (Jena Biosciences) PCR machine was used with the following program: 95°C for 1 min, then 40 cycles of 95°C for 15 s and 60°C for 30 s, followed by melting curve analysis with incremental temperature increase from 60°C to 100°C. The expression levels of amplified genes were normalized to the mean expression of *GAPDH* using the ΔCt method. All primers used are listed in **Table S5**.

### RNA sequencing and analysis

Total cell RNA was TRIzol-extracted from control and vU1-KO hiPSCs as per manufacturer’s instructions (Invitrogen), reverse-transcribed using random primers and ribodepleted before being sequenced on a Novaseq6000 platform (Illumina). Paired-end reads from each replicate were mapped to the reference human genome (build hg38) using STAR v.2.7.3a with default settings^59^. Mapped reads were quantified using htseq-count2 v.0.12.4^60^ and normalized via the *RUVs* function of the RUVseq package^61^. For differential expression analysis, vU1-KO and control were compared using DEseq2 v.1.34.0 and the default Wald test^62^. Genes with an adjusted *P*-value <0.05 and fold change (log_2_) >|0.6| were considered significantly differentially expressed. Functional enrichment analysis of gene sets was performed using Metascape^63^ or Gene Set Enrichment Analysis (GSEA; https://www.gsea-msigdb.org/).

### Splicing analyses

Individual RNA splicing events were detected in RNA-seq data, quantified, and compared between control and vU1-KO cells using Whippet^34^ with a probability cutoff of >0.9 and a |*Δ*psi| of >0.2. Functional changes to mRNA isoforms were detected, annotated, and quantified via *IsoformSwitchAnalyzeR* with default parameters^31^. Recursive splicing sites (RSSs) were first mapped using a custom in-house script to identify hybrid RNA-seq connecting the 3’ end of a given exon with downstream sequences in the following intron^49^. Next, RSSs were further filtered and quantified as previously described^56^, but considering all YAG|NN sequences (instead of AG|GT ones as in the original algorithm) and filtering out any RS events that were also detected in hiPSC poly(A)-enriched RNA-seq data. The number of reads supporting each RS event that fulfills all aforementioned criteria were normalized to all reads connecting the exons flaking the intron containing the RSS. A Student’s t-test with a *P*-value threshold of <0.05 was used when comparing conditions. The same approach was used in quantifying 3’ UTR usage levels, where the number of reads that map to the gene’s last exon, while 3’ UTR splicing was quantified using RNASeq3USP (https://github.com/christear/RNASeq3USP) and the same statistical comparison as above.

### vU1 RNA co-precipitation and analysis

For the co-precipitation of RNA interacting with vU1 sRNAs, we used 25 pmoles of a biotinylated 2’-O-Methyl RNA antisense oligonucleotide targeting vU1.3 or vU1.8 and ∼2 μg of iPSC total nuclear RNA. In brief, hiPSCs grown in 10-cm plates were crosslinked with psoralen as follows: AMT was first diluted in water at 1 mg/ml and then an equal volume of 2x PBS was added to produce a final concentration of 0.5 mg/ml, which was kept chilled on ice and in the dark at all times. Cells were washed once in 20 ml of ice-cold 1x PBS and the gently scraped into a 10-ml tube, centrifuged at 420 x g for 5 min at 4°C, and collected as a pellet. Following isolation of nuclei on ‘sucrose cushion’, hiPSC nuclei were resuspended in 4 ml of ice-cold AMT solution (or in ice-cold PBS alone that serves as the non-crosslinked control) and incubated on ice for 15 min, before being transferred to pre-chilled 10-cm cell culture dishes. The dishes, kept on ice, were placed under a 35 nm UV bulb in a Stratalinker 2400 (Stratagene) ∼3-4 cm from the light source for a total of 7 min at maximum power (while mixing gently every 2 min). Irradiated nuclei are then transferred to chilled tubes and spinned at 330 x g for 4 min, before pellets are treated with TRIzol (Invitrogen) to isolate total nuclear RNA and resuspended in 50 μl of water. The RNA yield is quantified (e.g., on a NanoDrop device) and for each 8 μg purified, a 50-μl reaction with 2.5 units of TURBO DNase (Invitrogen) is set up to degrade any genomic DNA contaminants and incubated at 37°C for 20 min. RNA in each reaction is re-purified using the miRNeasy kit (Qiagen) and eluted in 30 μl of water. Finally, 2.5 pmol of biotinylated 2’O-Methyl vU1-antisense oligonucleotide is added to 2 μg of nuclear RNA (for reference, for U1 snRNAs 20 pmoles are used) after denaturation of the oligonucleotide at 85 °C for 3 min and transferring to ice. The mixture is supplemented with 300 μl of pre-warmed hybridization buffer with LiCl, transferred to 37°C and incubated under shaking at 1200 rpm for 2 h. The crosslinked vU1:RNA complexes are pulled down using 20 μl of Streptavidin C1 magnetic beads per reaction, which has first been spun at 12,000 x g at 4°C to pellet debris. The beads and supernatants are incubated under shaking at 1,200 rpm, and then separated using a magnetic tube rack. The beads are last washed 3x in 250 μl of Low Stringency Wash Buffer at 37°C for 3 min/wash, and 3x in Low Stringency Wash Buffer the same way, before direct immersion in TRIzol to purify the co-precipitated RNAs.

vU1 pulldown targets were catalogued using RNA-seq. Due to low RNA yields, the eluates from three independent co-precipitations were pulled together and reversed transcribed using the SMART-Seq Total RNA Pico Input with UMIs kit (ZapR Mammalian; TAKARA Bio, 634354). The resulting libraries retain strand information and incorporate unique molecular identifiers (UMIs) during the reverse-transcription step to correct for PCR biases and assist transcript quantification. Following sequencing on a NovaSeq6000 platform (Illumina), vU1 pulldown targets were quantified with NOISeq^64^.

For the co-precipitation of proteins associated with vU1s, ∼50×10^6^ hiPSCs were used for isolating nuclei, resuspended in 20 mM Hepes (pH7.9), 100 mM KCl, 0.2 mM EDTA, 20% Glycerol, 0.5 mM PMSF plus 0.5 mM DTT, aliquoted to 10^7^ nuclei per reaction in the same without glycerol and with KCL and MgCl_2_ concentrations adjusted to 250 mM and 1.5 mM, respectively, and pelleted at 12,000 x g for 15 min at 4°C to remove any debris, before 5 pmoles of biotinylated 2’-O-Methyl vU1-antisense oligonucleotide per every 10^7^ nuclei (for reference, 30 pmoles are used for vU1 enrichment) and left under rotation at 4°C overnight. In parallel, 20 μl of Streptavidin Dynabeads are blocked in 1 ml of 20 mM Hepes pH 7.9, 150 mM KCl, 1.5 mM MgCl_2_, 0.1% NP-40, 50 μl BSA (from a 10 mg/ml stock) and 2.5 μl Glycogen (from a 20 mg/ml stock) at 4°C overnight. Next day, 10 μl of pre-blocked Dynabeads are added to the nuclear extract-antisense oligonucleotide mixture for 2.5 h at 4°C under end-over-end rotation, separated using a magnetic rack and collected, and then the rest 10 μl of beads are added and incubated for another 2.5 h. Beads from the back-to-back elution steps are pooled together, washed 4x in 20 mM Hepes (pH 7.9), 150 mM KCl, 1.5 mM MgCl_2_, 0.1% NP-40 for 10 min at 4°C under end-over-end rotation per wash, then washed 3x in ice-cold PBS for 5 min per wash, before eluting complexes associated to the beads in 50 μl elution buffer (2 M urea, 50 mM Tris pH 7.5, 1 mM DTT) supplemented by 50 ng trypsin for 30 min at room temperature with gentle agitation. Following addition of 50 μl digestion buffer (2 M Urea in 50 mM Tris pH 7.5, 5 mM chloroacetamide) and incubation for another 30 min at room temperature, an additional 50 μl of elution buffer supplemented with 50 ng of LysC and 100 ng of trypsin were added to each reaction and allowed to be digested overnight at room temperature. Finally, the digestion was stopped by adding 1 μl trifluoroacetic acid, the reactions were split in half, benzonase treated, purified on C18 stage tips, and all replicates were analyzed on a Q-Exactive platform (Thermo Fisher). MS results are listed in **Table S6**.

### Statistical analyses

All statistical tests (Wilcoxon, hypergeometric, t-test) were performed in R, except the Fisher’s exact test that was performed via the GraphPad online interface (https://www.graphpad.com/quickcalcs/contingency1/). Unless stated otherwise, results were deemed significant if they produced a P-value <0.01.

## Data availability

All RNA sequencing data generated as part of this study were deposited to the NCBI Gene Expression Omnibus repository and can be found under the accession number GSE305587 (reviewer access token: *inkdwmgavnilbod*).

## Acknowledgements

We thank Maria-Patapia Zafeiriou and Leo Kurian for their feedback, and members of the Papantonis lab for discussions. This study was supported by the German Research Foundation (DFG) via the Priority Program SPP1935 (313408820 awarded to A.P.), by the Lower Saxony Ministry for Research and Culture (MWK) via the SPRUNG program (76211-1267/2023 awarded to A.P.), as well as by the Scientific Core Facility for Cell Sorting of the University Medical Center Göttingen (DFG Large Equipment Project No. 442249343, LSR Fortessa X-20, BD). Y.Z. was further supported by the IMPRS Molecular Biology program.

## Author contributions

Y.Z. performed all computational analyses; K.S. and performed experiments together with A.M.; L.C. generated the vU1.8-KO hiPSC lines; M.N. performed GARLIC analyses; V.K. and C.F. performed differentiation experiments; A.P. conceived the project; Y.Z. and A.P. wrote the manuscript with input from all co-authors.

## Competing interests

The authors have no conflict of interest to declare.

## Supplementary Data

This Supplement contains **Figs S1-S4** and **Tables S1-S6**.

**Fig S1.**
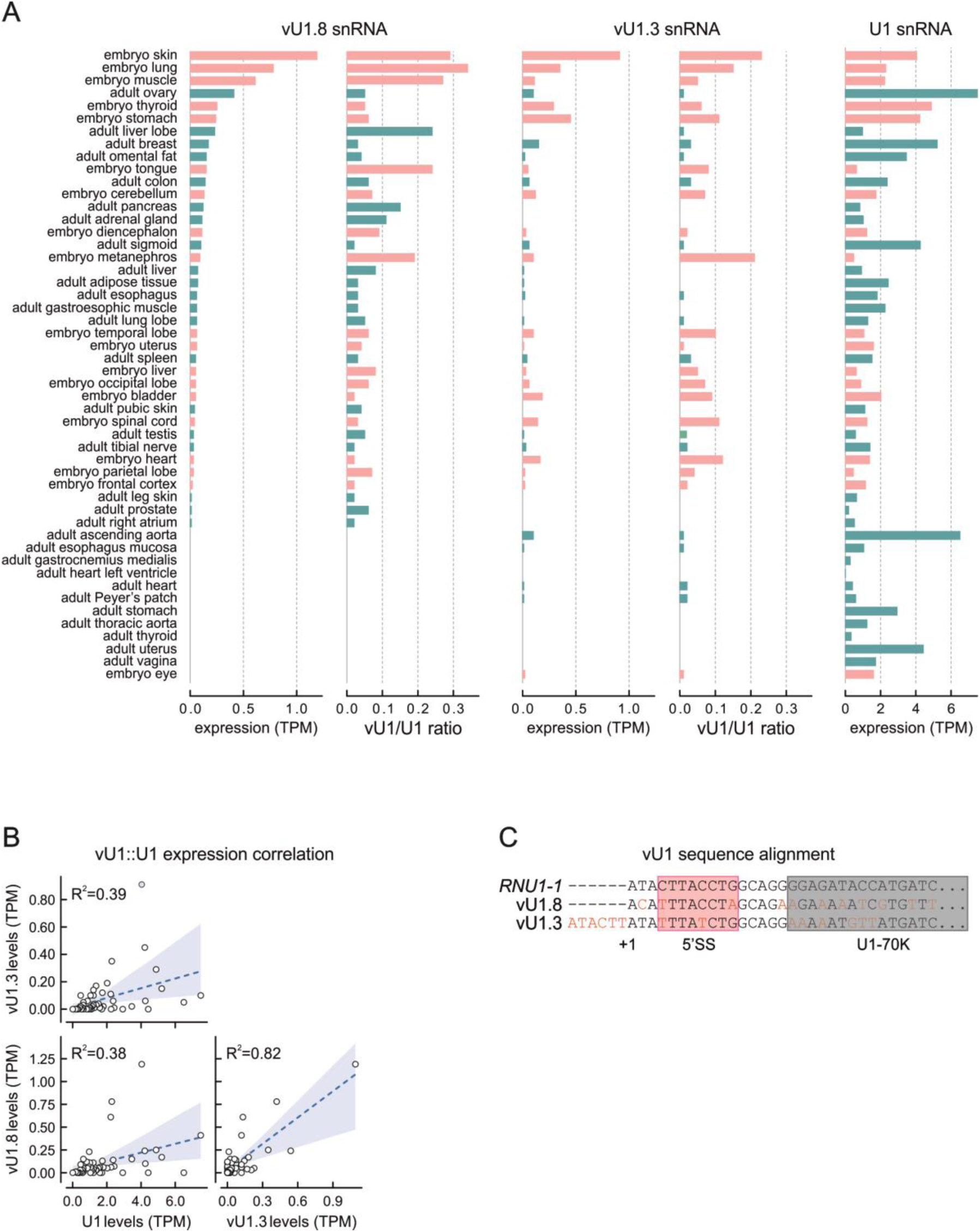
Expression of variant U1 snRNAs across human tissues. **a**, Bar plots showing the expression levels of vU1.3, vU1.8, and canonical U1 snRNA in embryonic (pink) and adult human tissues (green) from publicly-available RNA-seq data. **b**, Plots showing correlations of variant and canonical U1 snRNA levels from the data in panel A. **c**, Alignment of the vU1.3, vU1.8, and canonical U1 sequences around the region that interacts with donor splice sites on pre-mRNAs (pink).

**Fig S2.**
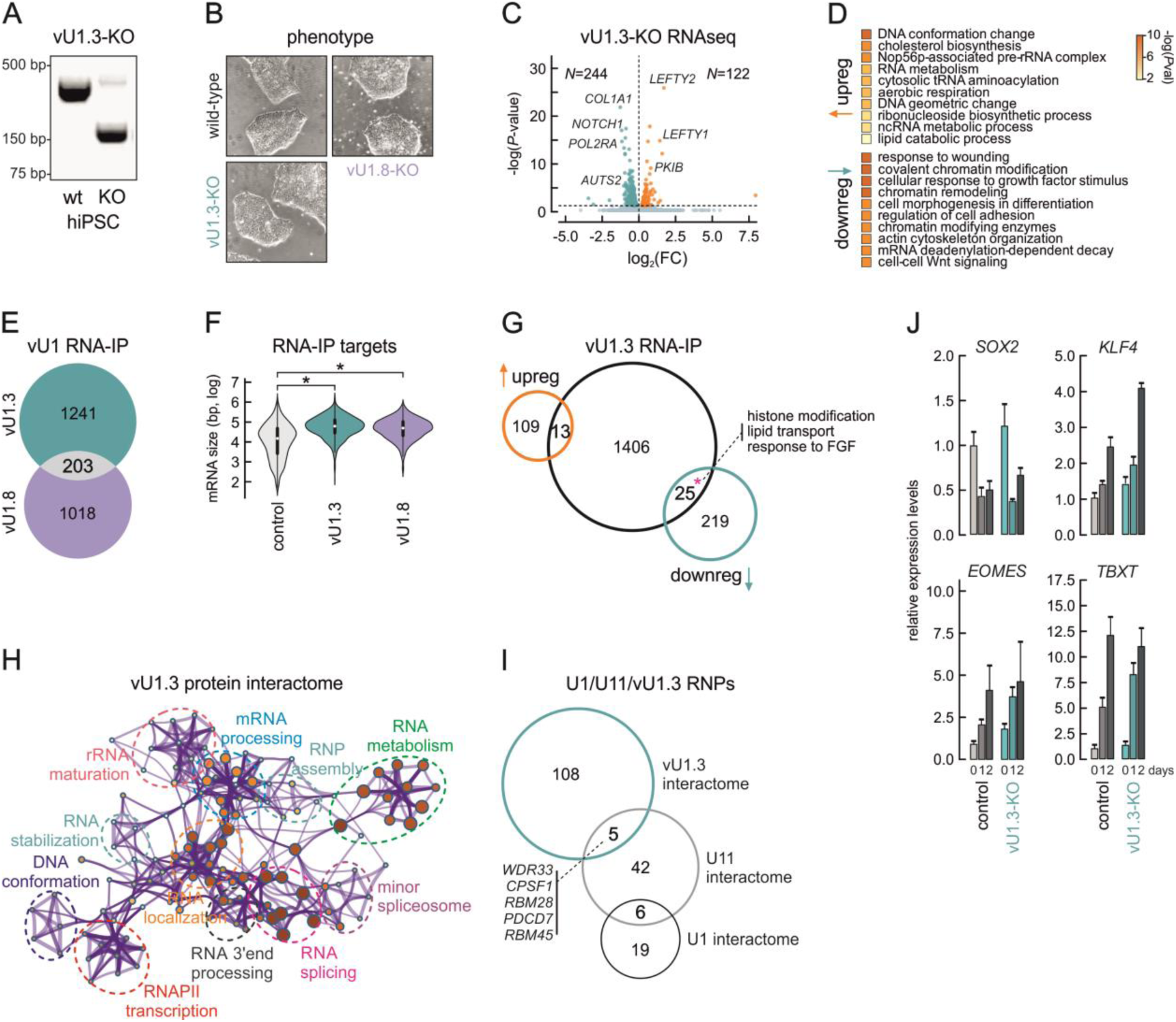
Effects of vU1-3 knockout in hiPS cells. **a**, Electrophoretic profiles of PCR products corresponding to the wild-type (wt) and knocked-out *RNU1-3* locus (vU1.3-KO). Molecular weight marker positions are indicated. **b**, Representative bright field images of wild-type, vU1.3-KO, and vU1.8-KO hiPSC colonies. **c**, Volcano plot showing differentially up-(orange) and downregulated genes (green) in vU1.3-KO hiPSCs given a *P*_adj_ cutoff of <0.05. **d**, Plot showing the top GO terms associated with vU1.3-KO up-/downregulated genes from panel c. **e**, Venn diagram showing the overlap of transcripts co-purified with vU1.3 (green) and vU1.8 (purple) in RNA-IP experiments. **f**, Violin plots showing the distribution of lengths for transcripts co-purified with vU1.3 (green) and vU1.8 (purple) compared to control ones from vU1.3-/1.8-KO hiPSCs. **g**, Venn diagram showing the overlap of differentially expressed genes from panel c with transcripts co-purified with vU1.3. **h**, Network representation of GO terms enriched for the proteins interacting with vU1.3. **i**, Venn diagram showing the overlap of proteins co-purified with vU1.3 with the core components of canonical U1 and U11. The proteins shared by the vU1.3 and U11 interactomes are shown. **j**, Bar plots showing changes (relative to day 0 ±S.D.) in expression levels of the indicated genes for control (ctrl) and vU1.8-KO hiPSCs at 1 or 2 days of differentiation.

**Fig S3.**
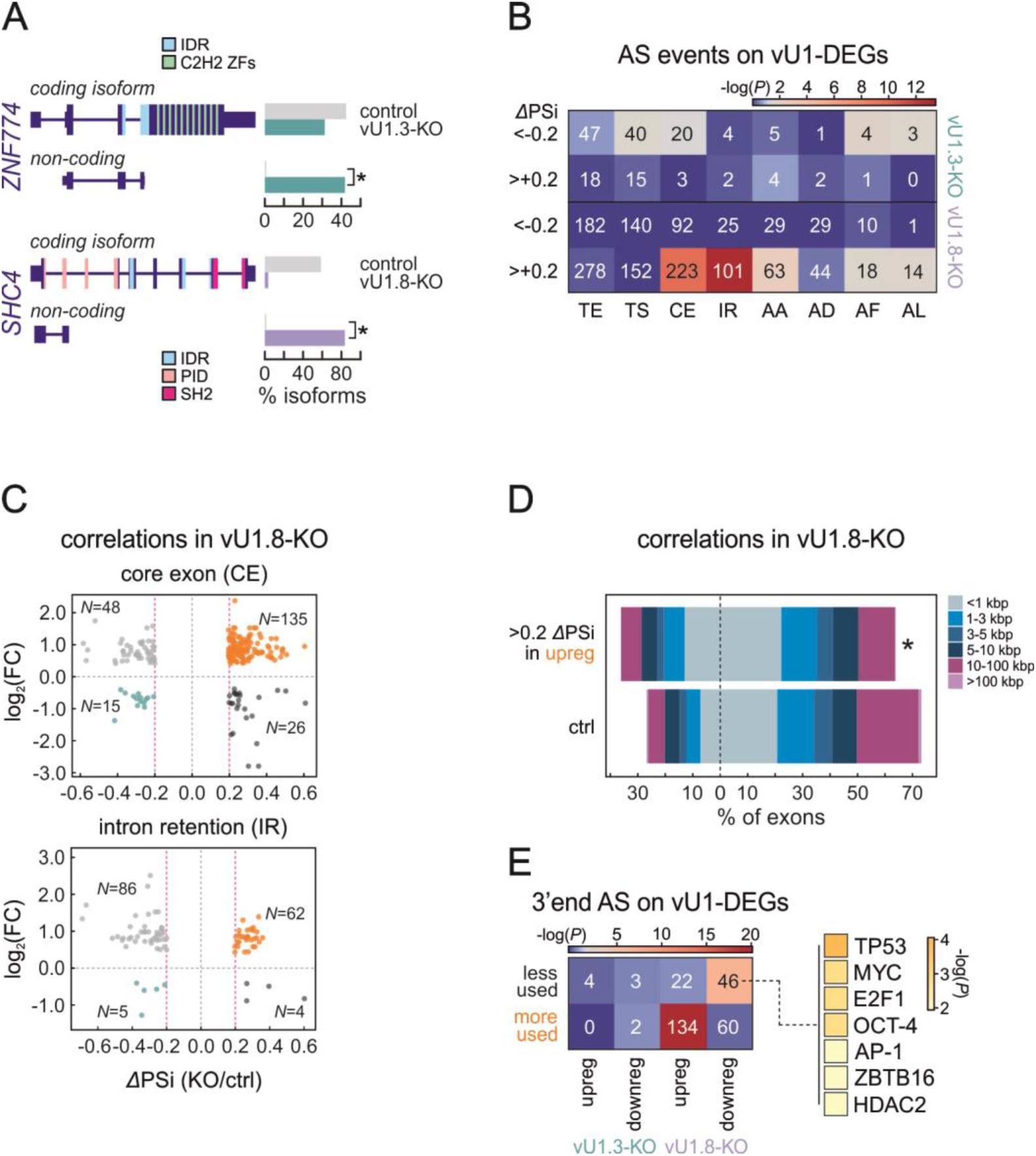
Alternative splicing effects upon vU1 knockout in hiPSCs. **a**, Changes in the levels of a coding (top) and non-coding *KLF5* isoform (bottom) in control (grey) and vU1.8-KO cells (purple). **b**, Heat map showing the number of events per AS type for differentially expressed genes (DEGs) in vU1.3-(top) or vU1.8-KO hiPSCs (bottom). The color scale represents statistical overrepresentation (-log*P*) in each case. **c**, Plot showing correlation between gene expression (log_2_FC) and splicing changes (*Δ*PSi) of vU1.8-associated transcripts for core exons (CE) or intron retention evens (IR). **d**, Plot showing the percent of exons located at increasing distances from TSSs showing differential usage in vU1.8-KO hiPSCs. **e**, Left: As in panel b, but for alternative 3’ end usage in each knockout line. Right: Enrichment for genes regulated by the indicated transcription factors (based on TTRUST^65^).

**Fig S4.**
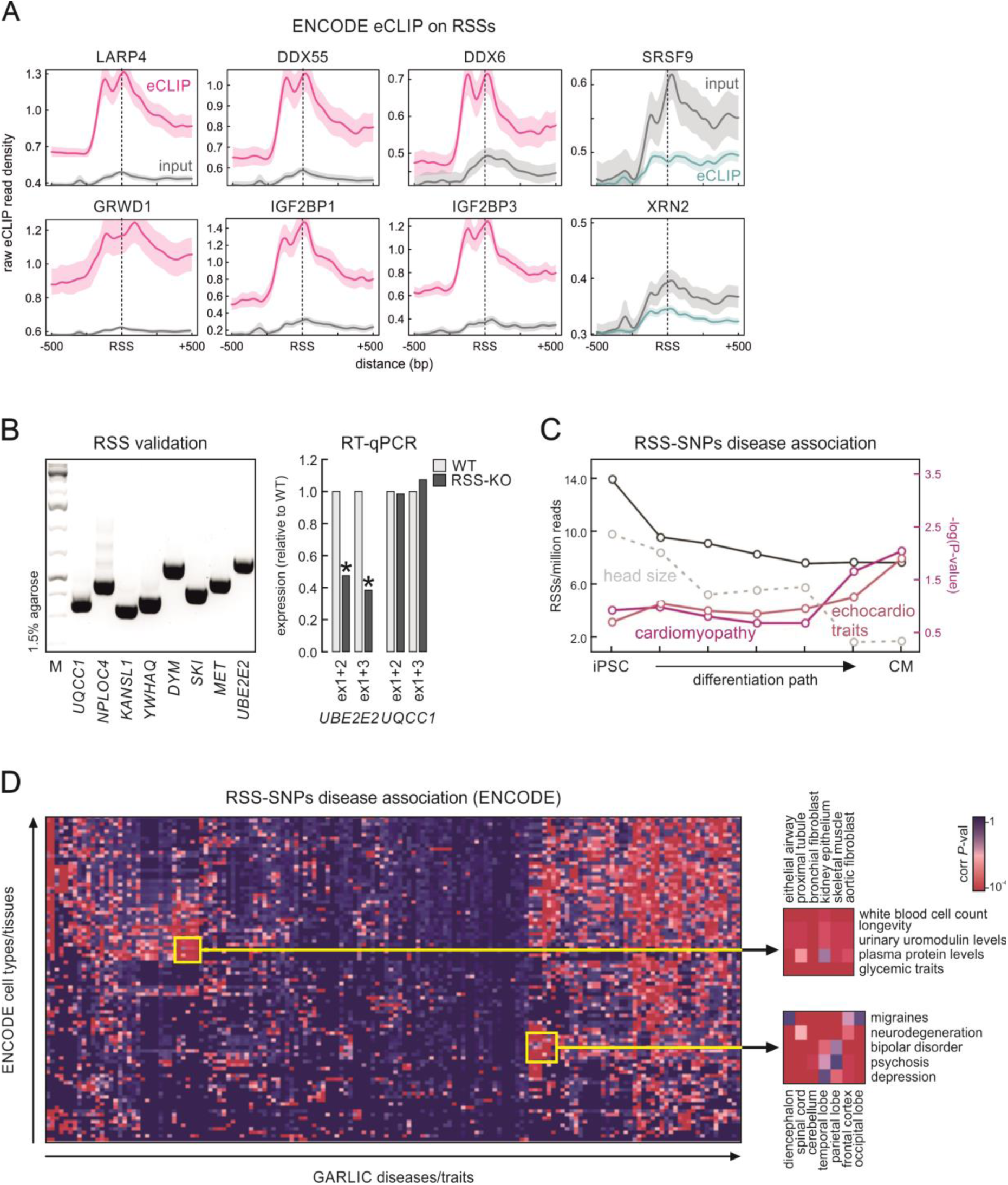
Features associated with recursive splicing sites. **a**, Plots showing mean eCLIP signal enrichment over RSSs for six RNA-binding proteins^53^. The non-enrichment of SRSF9 and XRN2 provide a negative control. **b**, Left: Electrophoresis profile of PCR-validate RSSs in eight different gene loci. Right: Bar plots showing changes in the mRNA levels of two exemplary genes upon RSS-knockout. **c**, Graph showing the number of RSSs discovered per million RNA-seq reads (black line) along the differentiation of hiPSCs into cardiomyocytes (RNA-seq data from ref. ^55^), and the enrichment for SNPs associated with cardiac disease (-log*P*; magenta/pink lines). Enrichment for SNPs associated with head size (dotted line) provides a control. **d**, Heatmap showing the statistical association between RSSs in ENCODE cell/tissue types and SNPs linked to diseases and traits in the GARLIC database^54^.

**Table S1.**
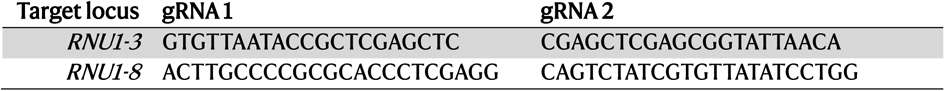
gRNAs used for the knockout of variant *RNU1* loci.

**Table S2.** List of differentially expressed genes in vU1.3- or vU1.8-KO hiPSCs (.xlsx file).

**Table S3.** Catalogue of transcripts interacting with vU1.3 or vU1.8 in RNA-IPs (.xlsx file).

**Table S4.** Alternative splicing events in vU1.3- or vU1.8-KO in hiPSCs (.xlsx file).

**Table S5.**
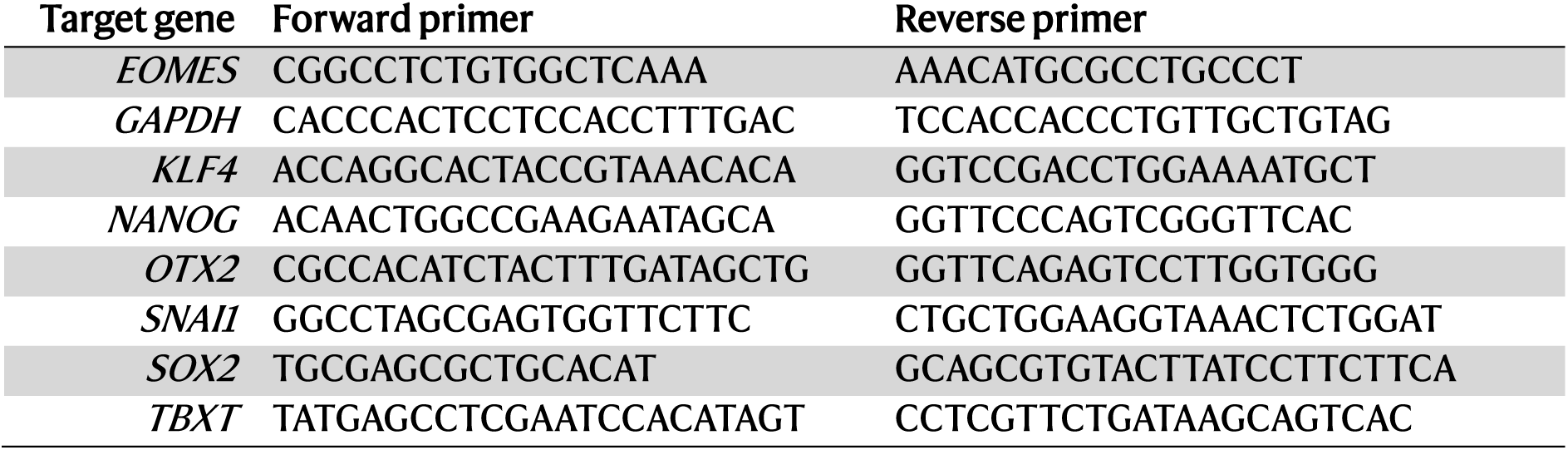
List of primers used in RT-qPCR.

**Table S6.** Catalogue of proteins interacting with vU1.3 in RNA-IPs (.xlsx file).

## Notes

### Competing Interest Statement

The authors have declared no competing interest.

